# OncoPhase: Quantification of somatic mutation cellular prevalence using phase information

**DOI:** 10.1101/046631

**Authors:** Donatien Chedom-Fotso, Ahmed Ashour Ahmed, Christopher Yau

## Abstract

The impact of evolutionary processes in cancer and its implications for drug response, biomarker validation and clinical outcome requires careful consideration of the evolving mutational landscape of the cancer. Genome sequencing allows us to identify mutations but the prevalence of those mutations in heterogeneous tumours must be inferred. We describe a method that we call OncoPhase to compute the prevalence of somatic point mutations from genome sequencing analysis of heterogeneous tumours that combines information from nearby phased germline variants. We show using simulations that the use of phased germline information can give improved prevalence estimates over the use of somatic variants only.

## 1 Introduction

Cancer exhibits extensive intra-tumour heterogeneity with multiple sub-populations of tumour cells containing both common and private somatic mutations. Within- and between-patient tumour heterogeneity is now seen as one of the major obstacles in precision medicine and the development of effective therapy strategies. Recent advances in genome sequencing technologies have enabled routine targeted or whole genome sequencing of tumour samples allowing the somatic landscape of a tumour to be surveyed but the existence of tumour heterogeneity makes it challenging to initially detect the existence of somatic mutations but also to determine the percentage of cancer cells harbouring those mutations (the prevalence). As the ability of cancers to evolve is increasingly recognised as an important factor in therapeutic success or failure [1–4], it is important to have accurate estimation of the prevalence of mutations to understand the evolution of the disease. The latter problem has recently given rise to a plethora of statistical approaches to reconstruct the underlying subclonal architecture of a tumour from sequencing of tumour samples containing mixed cell populations [5–10].

State-of-the-art methods for subclonal architecture reconstruction use statistical models that describe latent unobserved mutational profiles for an unknown number of tumour cell subpopulations which exist in different proportions in any given tumour sample. The inference problem is to infer the properties and prevalence of these subclones given the sequencing data which only gives an aggregated count of the number of variant (and non-variant) reads across all the cell populations in the tumour sample. One popular approach is to cluster the observed variant allele frequencies (VAFs) of each single nucleotide variant (SNV). The variant allele frequency is the ratio of the number of variant reads to the total number of reads covering that genomic locus. Each cluster of VAF therefore corresponds to one lineage in the evolutionary tree describing the sequence of mutational events affecting the tumour. This approach can be taken one step further by actually inferring the phylogenetic relationships themselves [6,9].

Current mainstream high-throughput sequencing typically adopts a short-read approach that sequences many short DNA fragments (30-300bp) in a highly multiplexed, parallel fashion enabling many times coverage of the whole genome. These short fragments are reassembled to provide sequential genomic coverage information by aligning to reference sequences. The result is that *phase* information is lost during the sequencing process precluding the ability to directly observe if a sequence variant lies on the same chromosome as another. However, recent adaptations of existing sequencing technologies and emerging new technologies (single molecule and nanopore techniques) have enabled longer physical reads to be obtained spanning 10s of kilobases or synthetic reads that potentially span many megabase regions.

In this paper we describe a methodology to quantify the cellular prevalence of single nucleotide variants (SNVs) using phase information. Our specific focus is on the accurate estimation of the prevalence of *particular* SNVs as supposed to full subclonal decomposition that is, we want to be able to provide to the answer “what proportion of cancer cells have this somatic mutation?”. Complete subclonal decomposition approaches utilise information across multiple SNVs to infer the prevalence and mutational profile of each subclone and the phylogenetic relationships between these subclones. The space of possible subclonal architectures is exponentially large, multiple subclonal configurations maybe compatible with the observed data and the prevalence of any individual SNV can vary depending on the lineage/tree combination to which it is attached.

Instead of tackling this complexity, our methodology relies on the fact that, within a population of tumour cells, germline variants (single nucleotide polymorphisms or SNPs) are always present in all the cells but SNVs may not. For any SNV, if we consider a SNP phased to it then that SNV, whenever present, will always be on the same chromosome with the SNP. In the case when neither the germline or somatic variant are affected by somatic copy number alterations (SCNAs), the ratio between the counts of sequencing reads supporting the SNV and the count of sequencing reads supporting the SNP will give an estimation of the somatic mutation prevalence. In the presence of SCNAs, if the copy number change occurred before the mutational event then the SNV will be less abundant than the SNP. In contrast, if the SNV occurs before the SCNA than the abundance of SNVs will match the SNPs. We exploit these relationships to show that the estimate of prevalence for phased SNVs can be increased. Furthermore, we are able to resolve ambiguities using the phase information due to the unknown latent subclonal architecture that cannot be done by looking exclusively at SNVs.

Accurate estimation of individual somatic mutations is important in applications where the mutational status of specific genes may have utility in assessing the efficacy of targeted therapies or stratified approaches to treatment. The methods we developed are implemented in a software package called OncoPhase that is freely available (https://github.com/chedonat/OncoPhase). OncoPhase uses haplotype phase information to accurately compute mutational prevalence. In the following we describe the mathematical formulation underlying the methodology and then illustrate its application in both simulated and real data settings.

## 2 Methods

OncoPhase uses a combination of phased SNV and SNP allele-specific sequence read counts and local allele-specific copy numbers to determine the prevalence of the SNV. Figure 1 gives a schematic of the methodology.

**Figure 1.**
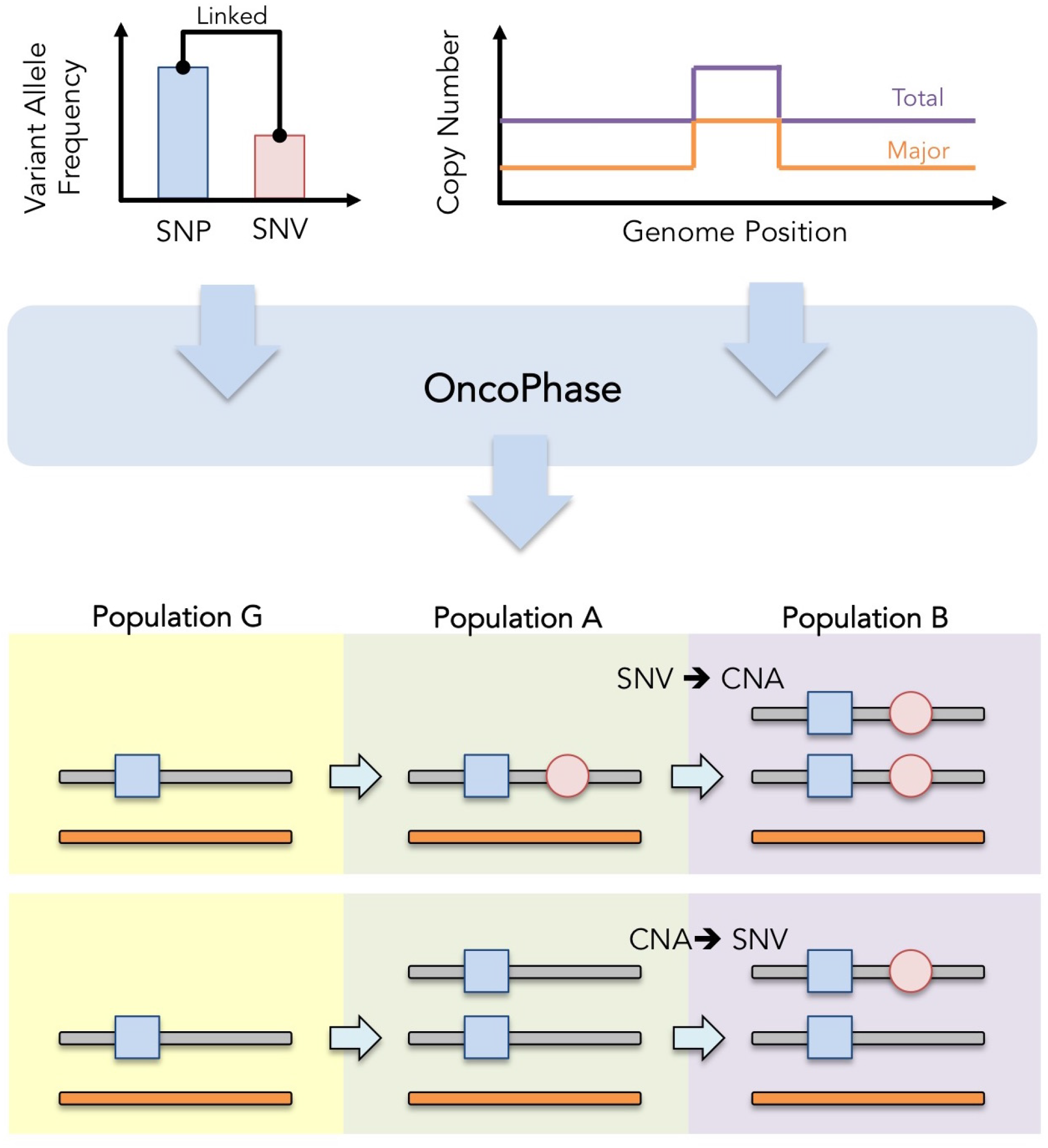
Workflow. OncoPhase takes as input phased read counts for SNP-SNV pairs (variant allele frequencies) and allele-specific copy number information and identifies (upto) three cell subpopulations (G, A, B).

### 2.1 Input specification

OncoPhase takes 7 (1 optional) input parameters:

1. (*λ_M_*,*μ_M_*) the allele counts for the somatic variant and reference allele at the mutation site *M*,
2. (*λ_G_*,*μ_G_*) the allele counts for the alternate and reference allele at the germline SNP *G* (where the alternate allele at *G* is in phased with the variant allele at *M*),
3. (*σ_G_*,*ρ_G_*) the allele-specific (alternate and reference) copy number at the locus *G* which can be obtained from appropriate copy number calling methods [11-13],
4. (Optional) *ψ* an estimate of the normal cell contamination

### 2.2 Prevalence computation

At each SNV, OncoPhase assumes the existence of three cellular sub-populations in the proportions *θ* = (*θ_G_, θ_A_, θ_B_*): (i) cells having a normal genotype with no mutations and no copy number alteration, (ii) cells harbouring either an SNV or a copy number alteration (but not both) and (iii) cells harbouring both the SNV and the copy number alteration. The cellular prevalence is therefore given by:

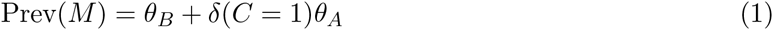

where *C* ∊ {0,1} is a latent indicator variable which has value 1 if the SNV occurs before the copy number alteration event and zero otherwise.

The variant allele frequency of the SNV *υ_M_* is given by:

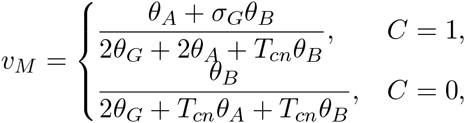

and the corresponding variant allele frequency for the phased SNP is given by:

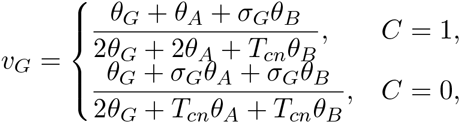

Given the latent variable *C* and the observed variant allele frequencies (*υ_M_*, *υ_G_*), the system of equations are linear in the parameters *θ* and can be solved using constrained linear solvers subject to the constraints *θ_G_* + *θ_A_* + *θ_B_* = 1 and 0 ≤ *θ_G_*, *θ_A_*, *θ_B_* ≤ 1. As *C* is unknown but only consists of two possibilities, the system can be solved for both possible configurations and the configuration giving the least residual error with respect to predicting the observed variant allele frequencies chosen.

Interestingly, in the absence of measurement noise, there is a closed-form solution for the prevalence of the mutation *M* given by

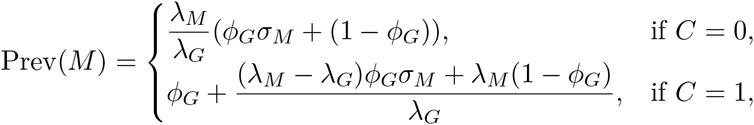

where 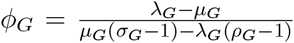. The existence of this closed form solution is instructive for understanding the utility of the use of phased information as we will detail in the following section.

In principle, the method can be used with any flanking SNP which is close to the SNV whether phased or not. However, in the absence of direct phasing information, it would also be necessary to solve the equations for the alternate phasing configurations.

## 3 Results

We conducted a simulation study to examine the utility of phasing information for estimating mutational prevalence. We simulated sequencing data, using a range of coverage levels (30, 60, 120 and 1000X), for the scenario depicted in Figure 2 which shows 6 sub-clonal cell types and their phylogenetic relationships. The simulation uses a variety of somatic point mutations and copy number changes. We treat the coverage levels as fixed and obtained the variant read counts for the SNVs and SNPs using Binomial sampling based on the expected variant allele frequency. We used 30 replicates to ensure robustness of our results to stochastic effects.

**Figure 2.**
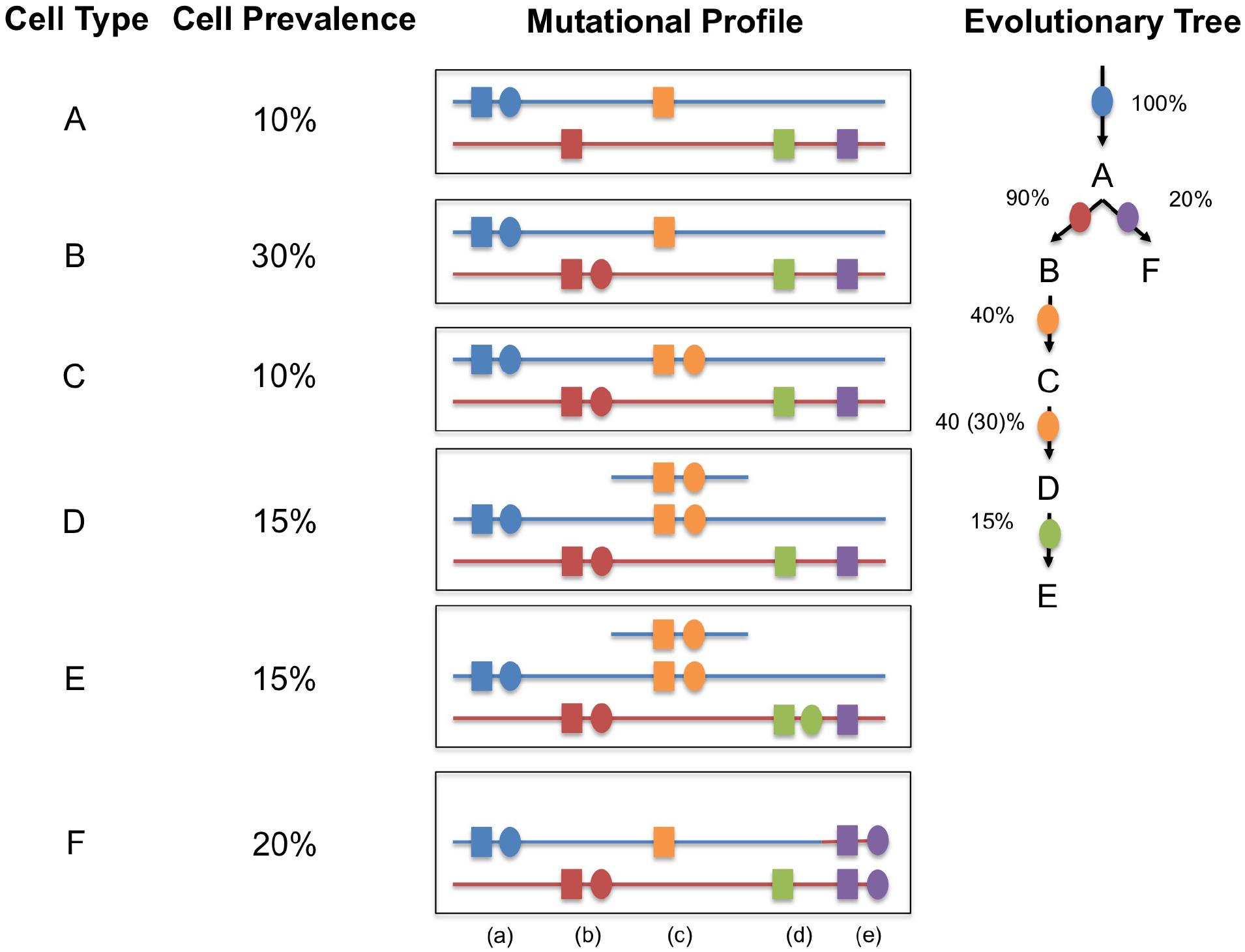
Simulation setup. The simulated data consists of six subclonal cell types containing a total of (a-e) 5 SNVs (o) and their respective local phased SNP (□). SNVs (c, e) are also affected by a duplication and copy-neutral LOH respectively.

We applied OncoPhase with and without the phased germline variant information to the simulated data set to compute the mutational prevalence for each of the 5 SNVs which are shown in Figure 3. For SNVs a, b and d, the use of phased germline information in OncoPhase was not advantageous, and it was possible to obtain good prevalence estimates using the SNV information alone. However, for SNVs c and e, the use of phasing information was vital. Without the phased germline information, the prevalence estimates were incorrect and did not improve with increased coverage, but OncoPhase was able to attain the correct values.

**Figure 3.**
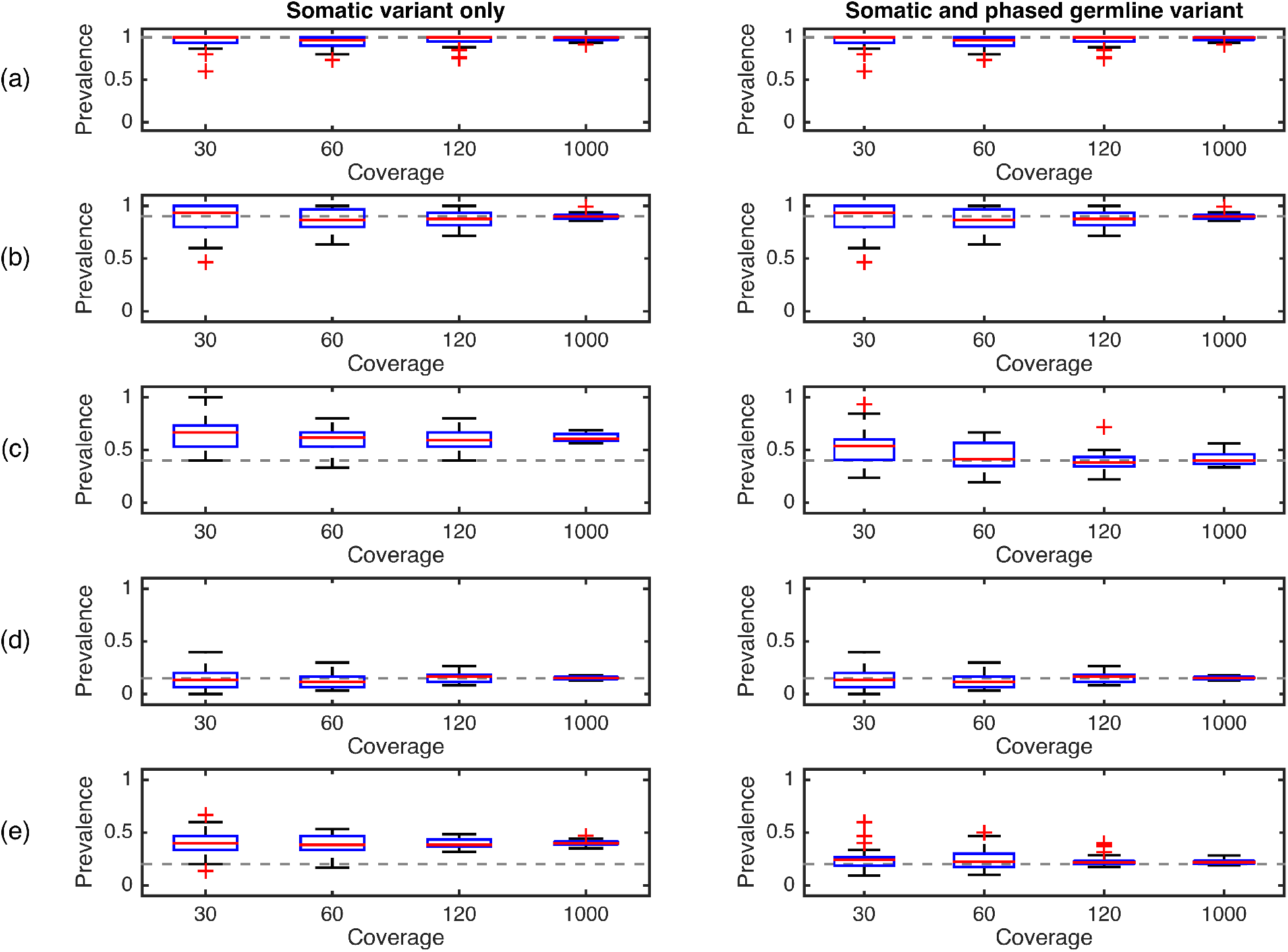
Simulation results. Estimated mutation prevalences using (i) somatic variants only and (ii) combining somatic variants with nearby phased germline variant information (OncoPhase). The rows (a-e) correspond to each of the five SNVs used in the simulation. The dashed line indicates the true mutational prevalence.

We next sought to further explore the properties of OncoPhase by examining simulations based on 12 specific scenarios shown in Figure 4. Table 1 gives illustrative noise-free input values based on these 12 cases that leads to the correct prevalence being estimated using the closed-form expression given previously. In the presence of noise, our simulations showed that OncoPhase was able to better estimate the mutational prevalences than an equivalent method based only on the SNV measurements (Figure 5). In cases 3, 4, 8, 9, 10, 11 and 12, it was not possible to obtain the correct prevalence levels using the SNV measurements only no matter the coverage level.

**Figure 4.**
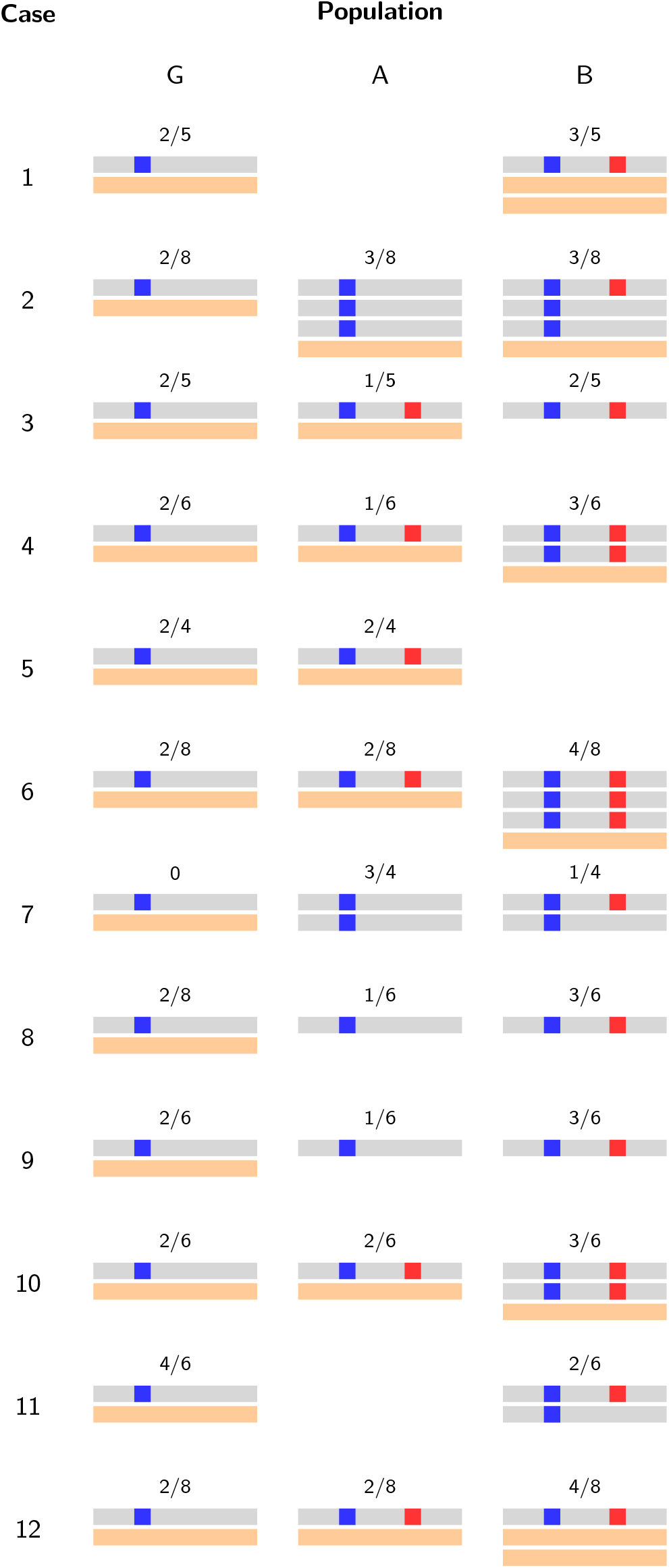
Further simulated examples. Twelve simulation scenarios. The colours blue denote a SNP and red denotes an SNV. The fractions above each chromosome denotes the proportions (*θ_G_, θ_A_, θ_B_*).

**Table 1.** Simulation parameters

| Case | $(\lambda_M, \lambda_G)$ | $(\mu_M, \mu_G)$ | $(\sigma_G, \rho_G)$ | $\phi_G$ | $C$ | $\text{Prev}(M)$ |
| --- | --- | --- | --- | --- | --- | --- |
| 1 | (14, 16) | (10, 8) | (3, 1) | 1/2 | 1 | 3/4 |
| 2 | (3, 20) | (25, 8) | (3, 1) | 3/4 | 0 | 3/8 |
| 3 | (3, 5) | (5, 3) | (1, 0) | 2/5 | 1 | 3/5 |
| 4 | (7, 9) | (8, 6) | (2, 1) | 1/2 | 1 | 2/3 |
| 5 | (2, 4) | (6, 4) | (1, 1) | 0 | 1 | 1/2 |
| 6 | (14, 16) | (10, 8) | (3, 1) | 1/2 | 1 | 3/4 |
| 7 | (1, 8) | (7, 0) | (2, 0) | 1 | 0 | 1/4 |
| 8 | (3, 6) | (5, 2) | (1, 0) | 2/3 | 0 | 1/2 |
| 9 | (6, 7) | (2, 1) | (2, 0) | 3/4 | 1 | 3/4 |
| 10 | (6, 8) | (8, 6) | (2, 1) | 1/3 | 1 | 2/3 |
| 11 | (4, 8) | (8, 4) | (2, 0) | 1/3 | 1 | 1/3 |
| 12 | (6, 8) | (14, 12) | (1, 2) | 1/2 | 1 | 3/4 |

**Figure 5.**
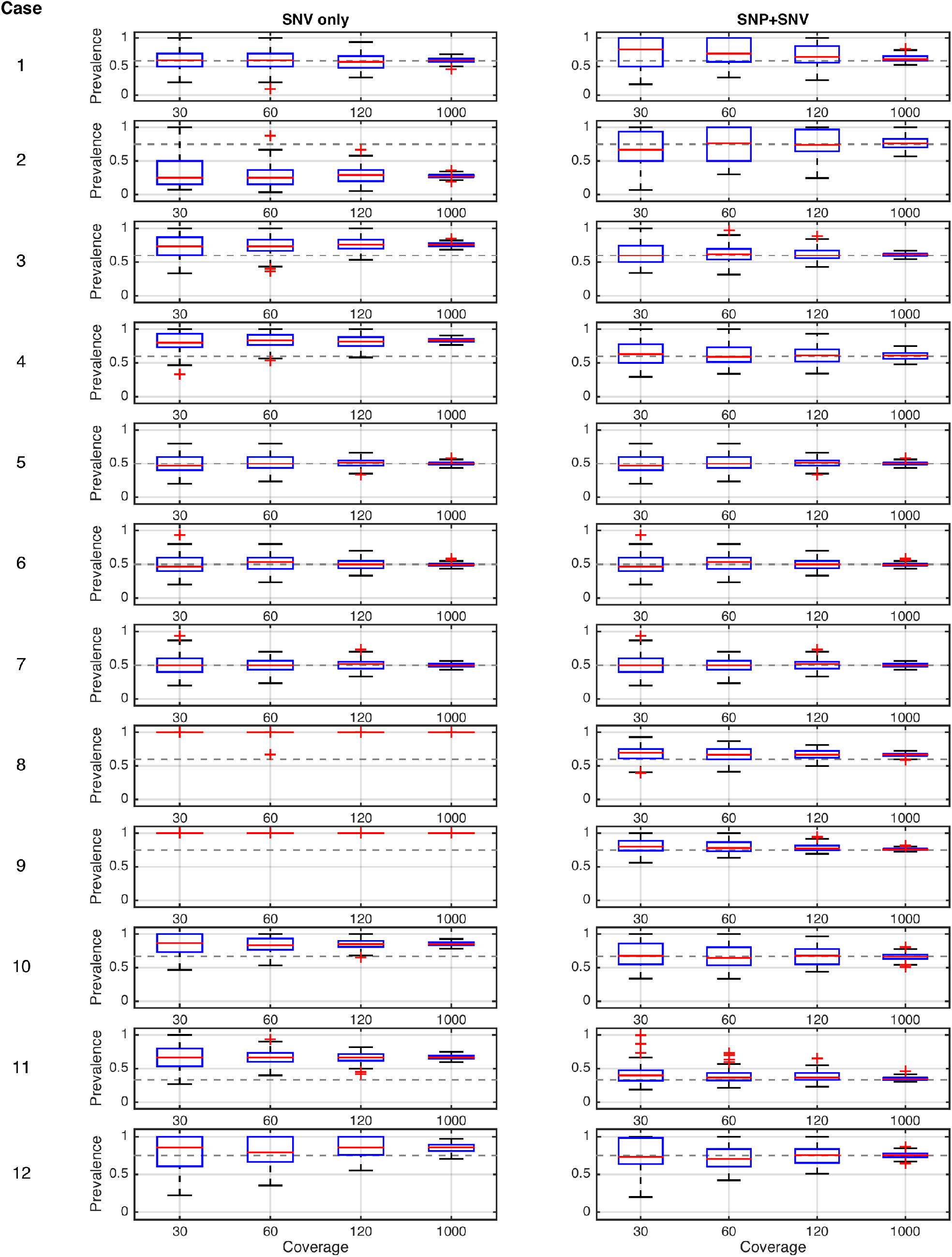
**Comparison of OncoPhase prevalence estimates for the twelve further simulated examples with and without phased germline information**.

## 4 Discussion

The accurate inference of mutational prevalence levels is critical for tracking subclonal dynamics in cancer. Full clonal architecture models allow for the simultaneous inference of subclonal cell types, mutational profiles and their phylogenetic relationships. These methods leverage information across multiple (potentially thousands) SNVs and copy number alterations across the whole genome but this results in less specificity for certain mutations as the objective is to obtain a good global model fit to the data.

Our method OncoPhase has been designed to focus on target mutations *only*. It leverages information from adjacent SNPs that can be phased with the somatic variant and combines the two sources of data to improve mutational prevalence estimates. The use of local phased SNPs also helps to resolve potential ambiguities that would confound estimation using the somatic variant data alone. Our methodology is timely with the emergence of sequencing methods that are capable of producing either actual physical long-reads [14,15] or synthetic long-reads [16–18] that gives phasing information.

## Acknowledgements

CY is supported by a Wellcome Trust Core Award Grant Number 090532/Z/09/Z and a UK Medical Research Council New Investigator Research Grant (Ref. No. MR/L001411/1). DCF, CY and AAA are supported by a Research Grant from the Ovarian Cancer Action Charity. AAA is supported by the Medical Research Council and the Oxford Biomedical Research Centre, the National Institute of Health Research.

## Contributions

AAA and CY conceived the study. DCF and CY developed the methods and wrote the software. All authors contributed to the manuscript.

